# Modulation of Stemness and Differentiation Regulators by Valproic Acid in Medulloblastoma Neurospheres

**DOI:** 10.1101/2024.09.23.614476

**Authors:** Natália Hogetop Freire, Alice Laschuk Herlinger, Julia Vanini, Matheus Dalmolin, Marcelo A. C. Fernandes, Carolina Nör, Vijay Ramaswamy, Caroline Brunetto de Farias, André Tesainer Brunetto, Algemir Lunardi Brunetto, Lauro José Gregianin, Mariane da Cunha Jaeger, Michael D. Taylor, Rafael Roesler

## Abstract

Changes in epigenetic processes such as histone acetylation are proposed as key events influencing cancer cell function and the initiation and progression of pediatric brain tumors. Valproic acid (VPA) is an antiepileptic drug that acts partially by inhibiting histone deacetylases (HDACs) and could be repurposed as an epigenetic anticancer therapy. Here, we show that VPA reduced medulloblastoma (MB) cell viability and led to cell cycle arrest. These effects were accompanied by enhanced H3K9 histone acetylation (H3K9ac) and decreased expression of the *MYC* oncogene. VPA impaired the expansion of MB neurospheres enriched in stemness markers and reduced *MYC* while increasing *TP53* expression in these neurospheres. In addition, VPA induced morphological changes consistent with neuronal differentiation and the increased expression of differentiation marker genes *TUBB3* and *ENO2*. The expression of stemness genes *SOX2*, *NES*, and *PRTG* was differentially affected by VPA in MB cells with different *TP53* status. VPA increased H3K9 occupancy of the promoter region of *TP53*. Among the genes regulated by VPA, the stemness regulators *MYC* and *NES* showed an association with patient survival in specific MB subgroups. Our results indicate that VPA may exert antitumor effects in MB by influencing histone acetylation, which may result in the modulation of stemness, neuronal differentiation, and the expression of genes associated with patient prognosis in specific molecular subgroups. Importantly, the actions of VPA in MB cells and neurospheres include a reduction in the expression of *MYC* and an increase in *TP53*.

## 1. Introduction

Medulloblastoma (MB) is the most common malignant childhood brain tumor. MB arises from neural stem cells (NSCs) or cerebellar granule neuron precursors (GNPs) that undergo genetic and epigenetic alterations [1–4]. Genomic, epigenomic, and transcriptional analyses have shown that MB is a heterogeneous disease with a variety of molecular, clinical, and prognosis features. Current classification has established that MB comprises four distinct molecular subgroups: WNT-activated, SHH-activated, and non-WNT/non-SHH MB that includes group 3 and group 4 MB, which have a particularly poor prognosis [3–7]. Within the SHH subgroup, *TP53* status is crucial, with tumors harboring *TP53* mutations defining a particularly high-risk group of patients. *TP53* mutations are commonly associated with downstream changes including amplification of the *MYC* paralog *MYCN* [8]. *MYC* is an oncogene frequently amplified or overexpressed in many cancer types, resulting in increased cell proliferation [9].

Despite advancements in therapies in the past few decades, metastatic and recurrent MB tumors remain a challenge. Relapsed MB occurs in approximately 30% of patients and have high mortality rates [4,10]. A subpopulation of cells, known as cancer stem cells (CSCs), contributes to treatment resistance and metastasis occurrence [11–13]. CSCs display stem cell features such as self-renewal and differentiation potential. Neurospheres derived from MB tumors or cell lines express markers related to neural progenitors and stem cells such as CD133, SOX2, and BMI-1 proto-oncogene, polycomb ring finger (BMI-1) [14]. Moreover, CD133 positive (CD133+) MB cells are able to initiate tumors similar to those of the patient’s original ones when implanted in NOD-SCID mice [11,15], supporting the view that CSCs contribute to MB carcinogenesis.

Epigenetic mechanisms are major drivers in the establishment and maintenance of CSCs. Most common mutations found in cancer are structurally or functionally related to epigenetic regulators [16]. An altered epigenetic profile enables cellular reprogramming that contributes to the aberrant activation of stem cell pathways promoting the acquisition of uncontrolled self-renewal [17,18]. Moreover, the choice between an undifferentiated and differentiated state can be controlled by alterations in the epigenome [19]. Valproic acid (VPA) is an anticonvulsant drug that also acts as an epigenetic modulator capable of inhibiting histone deacetylases (HDACs). VPA specifically inhibits HDAC class I and class IIa (HDAC 1–5, 7), leading to enhanced histone acetylation [20]. HDAC inhibitors show therapeutic potential in experimental MB [21–28]. VPA was shown to be associated with changes in MB cell cycle progression, senescence, and apoptosis [29]. In glioblastoma CSCs, VPA downregulates the expression of stemness genes *Prominin 1* (*PROM1*, which encodes CD133), *NANOG*, and *POU Class 5 Homeobox 1* (*POU5F1*, also known as *OCT4*) and increases differentiation markers [30]. Hence, VPA may modulate the epigenome and contribute to CSC maintenance. Here, we explore the role of VPA in influencing genes related to differentiation and stemness maintenance in MB.

## 2. Materials and Methods

### 2.1. Cell Culture and Drug Treatment

Human MB D283 (ATCC^®^ HTB-185™) and Daoy (ATCC^®^ HTB-186™) cells were originally obtained from the American Type Culture Collection (ATCC, Rockville, ML, USA). D283 (*TP53* wild-type) and Daoy (*TP53*-mutated) cells are proposed to be representative of group 3 and SHH MB, respectively [31]. Cells were maintained in Dulbecco’s modified Eagle’s medium (DMEM low glucose, Gibco^®^, Thermo Fisher Scientific, Waltham, MA, USA) containing 10% (*v*/*v*) fetal bovine serum (FBS, Gibco), 1% (*v*/*v*) penicillin-streptomycin solution (10,000 U/mL, Gibco), and 0.1% (*v*/*v*) amphotericin B (250 μg/mL; Gibco). Cells were cultured at 37 °C in a humidified incubator under 5% CO_2_. Experiments were conducted in exponentially growing cell cultures. VPA (CAS 1069-66-5, Santa Cruz Biotechnology, Dallas, TX, USA) was dissolved in sterile water to a stock concentration of 0.3 M.

### 2.2. Cell Viability

MB cells were treated with VPA (0.5, 1.0, 2.5, 5.0, 10.0, or 20.0 mM) for 48 h. Cells were seeded at 3000 cells/well in 96-well plates and after VPA exposure, they were detached with trypsin-EDTA (Gibco) and counted in a Neubauer chamber with trypan for the viability measurement. Doses of VPA were chosen based on previous in vitro studies using cultured MB cell lines [29,32]. Experiments were conducted in three biological replicates. For half maximal inhibitory concentration (IC_50_) determination, cell viability data were fitted in a dose–response curve (GraphPad Prism v. 8.0).

### 2.3. Cell Cycle

To assess the cell cycle distribution, MB control and VPA-treated cells were detached, centrifuged, and washed with PBS twice. Cells were then resuspended in 50 μg/mL propidium iodide (Sigma-Aldrich, St. Louis, MO, USA) in 0.1% Triton X-100 in 0.1% sodium citrate solution and incubated on ice for 15 min. The cells were analyzed by flow cytometry (Attune^®^, Applied Biosystems, Thermo Fisher Scientific) and 20,000 events were collected per sample.

### 2.4. Neurosphere Formation

A neurosphere formation assay was used as a model to expand putative CSCs on the basis of previous studies [22,33]. To analyze the effects of VPA during neurosphere formation, MB cells were dissociated with trypsin-EDTA into a cell suspension and seeded at 500 cells/well in 24-well plates. Agarose solution (1%) was used to overcome cell adherence. Cells were cultured in a serum-free neurosphere-induction medium containing DMEM/F12 supplemented with 20 ng/mL epidermal growth factor (Sigma-Aldrich, St. Louis, MO, USA), 20 ng/mL basic fibroblast growth factor (Sigma-Aldrich), 2% B-27 supplement (Gibco), 0.5% N-2 supplement (Gibco, Life Technologies), 50 μg/mL bovine serum albumin (Sigma Aldrich), and antibiotics for 5 days as previously described [22]. Cells were monitored daily until neurosphere formation. To analyze the effects during neurosphere formation, VPA (1.0, 2.5, 5.0, 10.0, or 20.0 mM) was added on the first day of the neurosphere formation assay and the neurosphere size was measured after a period of 5 days. To verify VPA modulation after neurosphere formation, MB cells were dissociated with trypsin-EDTA into cell suspension and seeded at either 500 or 1000 cells/well in 24-well ultra-low attachment plates (Corning^®^) in serum-free neurosphere-induction medium. After 5 days, VPA was added at a final concentration corresponding to half the IC_50_ (D283, 2.3 mM; Daoy, 2.2 mM), and the neurosphere size and number were analyzed after 48 h. Images were taken using an inverted microscope at 5 X amplification and the neurosphere size was measured using ImageJ version 1.53 (National Institutes of Health, Bethesda, ML, USA). Experiments were conducted in three biological replicates.

### 2.5. Reverse Transcriptase Polymerase Chain Reaction (RT-qPCR)

Messenger RNA expression of target genes was analyzed by RT-qPCR. RNA was extracted from MB monolayer cells or neurospheres using the ReliaPrep™ RNA Miniprep System (Promega, Madison, WI, USA), in accordance with the manufacturer’s instructions and quantified in NanoDrop (Thermo Fisher Scientific). The cDNA was obtained using the GoScript Reverse System (Promega), also according to the manufacturer’s instructions. The mRNA expression levels of cyclin-dependent kinase inhibitor 1 (*CDKN1A*), *ENO2*, *NES*, *MYC*, *SOX2*, *TUBB3,* and protogenin (*PRTG*) were quantified using PowerUp SYBR Green Master Mix (Applied Biosystems, Thermo Fisher Scientific). Primers used for RT-qPCR amplification were designed according to the literature and are shown in Supplementary Table S1. Expression of actin beta (ACTB) was measured as the control.

### 2.6. Western Blot

MB control and VPA-treated cells were lysed with 1 X lysis buffer (Cell Lysis Buffer, Cell Signaling Technology, Danvers, MA, USA), and protein was quantified using the Bradford protein assay (Bio-Rad Laboratories, Hercules, CA, USA). For blotting, 40 µg of protein was separated by SDS-PAGE and transferred to a PVDF membrane. Membranes were blocked for 1 h (5% skim milk in TTBS) prior to overnight incubation at 4 °C with primary antibodies against p21 (1:200; Santa Cruz Biotechnology sc-6246) and β-actin (1:2000; Santa Cruz Biotechnology sc-47778) as the loading control. The incubation of primary antibodies was followed by incubation with the secondary antibody adequate to each primary antibody for 1 h. Chemiluminescence was detected using ECL Western Blotting Substrate (Pierce, Thermo Fisher Scientific), and images were obtained using iBright (Thermo Fisher Scientific). Immunodetection signals were analyzed using ImageJ (National Institutes of Health).

### 2.7. Immunofluorescence

Cells were seeded into coverslips treated with poly-L-lysine solution 0.01% (Sigma Aldrich) and exposed to VPA for 48 h, while neurospheres were moved to coverslips treated with poli-L-lysine solution 0.01% after VPA exposure. MB monolayer cells and neurospheres were washed with phosphate-buffered saline (PBS), fixed with methanol for 5 min at room temperature (RT), and washed 2 times with ice cold PBS. Coverslips were incubated for 30 min at RT in blocking solution (1% of bovine serum albumin (BSA), 0.1% Tween 20 in PBS) prior to incubation with primary antibodies against the histone H3 lysine 9 acetylated residue (H3K9ac) (1:3000; Abcam ab177177), histone H3 (1:250; Thermo Fisher Scientific MA3-049), and beta III tubulin (1:1000, Abcam ab18207) at 4 °C overnight. Coverslips were then rinsed three times with PBS and incubated with secondary fluorescent antibodies Alexa Fluor 488-conjugated goat anti-rabbit (1:1000; Abcam ab150077) and Alexa Fluor 594-conjugated anti-mouse (1:1000; Abcam ab150116) for 1 h at RT. Monolayer cells and neurospheres were then washed with PBS, and coverslips were mounted with Fluoroshield-containing DAPI (Sigma Aldrich) to counterstain the nuclei. Fluorescent monolayer cells and neurospheres were examined using a Leica microscope under 10 or 20 x magnification. Images of DAPI or H3K9ac were binarized, and the intensity of H3K9ac normalized to DAPI was measured using ImageJ for semi-quantitative analysis.

### 2.8. Chromatin Immunoprecipitation (ChIP)

D283 cells were seeded at a density of 1.5 × 10^6^ cells per flask in T75 culture flasks previously coated with 1% agarose. Cells were cultured in a serum-free neurosphere-induction medium (8 mL per flask). Fresh medium (8 mL) was added on the third day of neurosphere induction. On the fifth day, the same amount of fresh media was added containing VPA at a final concentration of 2.3 mM. Three independent experiments were performed in parallel, each with four replicate flasks, which were pooled (using an equal number of cells from each independent experiment) to achieve a total of 4 × 10^6^ cells, which were then used for the ChIP experiments. ChIP was performed using the Pierce Agarose ChIP Kit (Thermo Fisher Scientific), following the manufacturer’s instructions with modifications. Immediately after preparing the cell pools, crosslinking was performed by incubating cells with 1% formaldehyde for 10 min at room temperature. Cells were subsequently incubated with glycine for 5 min at room temperature. Cells were then washed twice with ice-cold PBS (5000× g, 3 min) and once with icecold PBS containing protease inhibitors (5000× g, 3 min). Following cellular and nuclear membrane lysis according to the manufacturer’s instructions, chromatin was fragmented with 6U of micrococcal nuclease per 2 × 10^6^ cells as per the manufacturer’s instructions. The IP was performed using the Anti-Histone H3 (acetyl K9) antibody—ChIP Grade (Abcam ab108123; 3 μg/ 2 × 10^6^ cells) overnight at 4 °C. Normal rabbit IgG (provided with the kit) was used as a negative control. The IP chromatin was recovered according to the manufacturer’s instructions and stored at −80 °C until qPCR. qPCR was performed using the PowerUp SYBR Green Master Mix (Thermo Fisher) and 4 μL of IP chromatin per reaction, using a QuantStudio3 Real-Time System (Thermo Fisher Scientific). Primers targeting the promoter region of *TP53* are shown in Supplementary Table S2. Experiments were performed in quadruplicate and the results are presented as % input.

### 2.9. Gene Expression in MB Tumors and Patient Survival

Analyses of the *MYC* and *NES* transcripts were performed in a previously described transcriptome dataset from patients with MB (Cavalli cohort, (GEO: GSE85218)) [34]. These two genes were chosen for analysis on the basis of the amount of information available in the dataset. We selected 612 MB tumor samples with associated patient survival information, classified as follows: 113 group 3 MB samples, 264 group 4 samples, 172 SHH MB samples, and 64 WNT tumors. Expression levels were normalized using the R2 Genomics Analysis and Visualization Platform (http://r2.amc.nl). Tumors were classified into the different molecular subgroups and subtypes according to data available in the dataset. A feature of the R2 platform, namely the Kaplan Scan (KaplanScan algorithm), where an optimum survival cut-off is established based on statistical testing, was used.

### 2.10. Statistics

Data are shown as the mean ± standard deviation (SD). Statistical analyses for cellular and molecular experiments were performed by an independent Student’s *T*-test when comparing two groups, or one-way analysis of variance (ANOVA) followed by Bonferroni post hoc tests for multiple comparisons. Experiments were replicated at least three times; for the ChIP-qPCR data, two-way ANOVA with the Sidak test was performed; *p* values under 0.05 indicated statistically significant differences. The GraphPad Prism 8 software (GraphPad Software, San Diego, CA, USA) was used for analyses.

For gene expression analysis in the tumor samples, patients were classified into high and low gene expression groups with the “Survminer” package with ‘minprop = 0.2’. To investigate differences between subgroups, we employed the Wilcoxon test and subsequently used the Dunn test. Statistical significance was assessed using the Holm-adjusted *p* value test. These analyses were conducted utilizing the ‘ggstatsplot’ package. Patient overall survival (OS) was analyzed using the “Survival” package. Patient OS was measured from the day of diagnosis until death or the date of last follow-up. OS was calculated using the Kaplan–Meier estimate with median values and log-rank statistics, with *p* < 0.05 indicating significant differences between groups.

## 3. Results

### 3.1. VPA Reduces MB Cell Viability

To evaluate the effects of VPA inhibition on MB cell viability, we exposed D283 and Daoy cells to different concentrations of VPA (0.5, 1.0, 2.5, 5.0, 10.0, or 20.0 mM) for 48 h, and viable cells were counted in a Neubauer chamber with trypan blue. Consistently with previous evidence [29], VPA reduced the cell number in a dose-dependent manner (Figure 1A), suggesting a decrease in cell viability. Estimated IC_50_ values showed that both cell lines presented similar sensitivities to VPA (2.3 mM for D283 and 2.2 mM for Daoy cells). In addition, an immunofluorescence assay was used to confirm the association in the VPA-induced impairment of viability with its capability to increase histone H3K9ac (Figure 1B).

**Figure 1.**
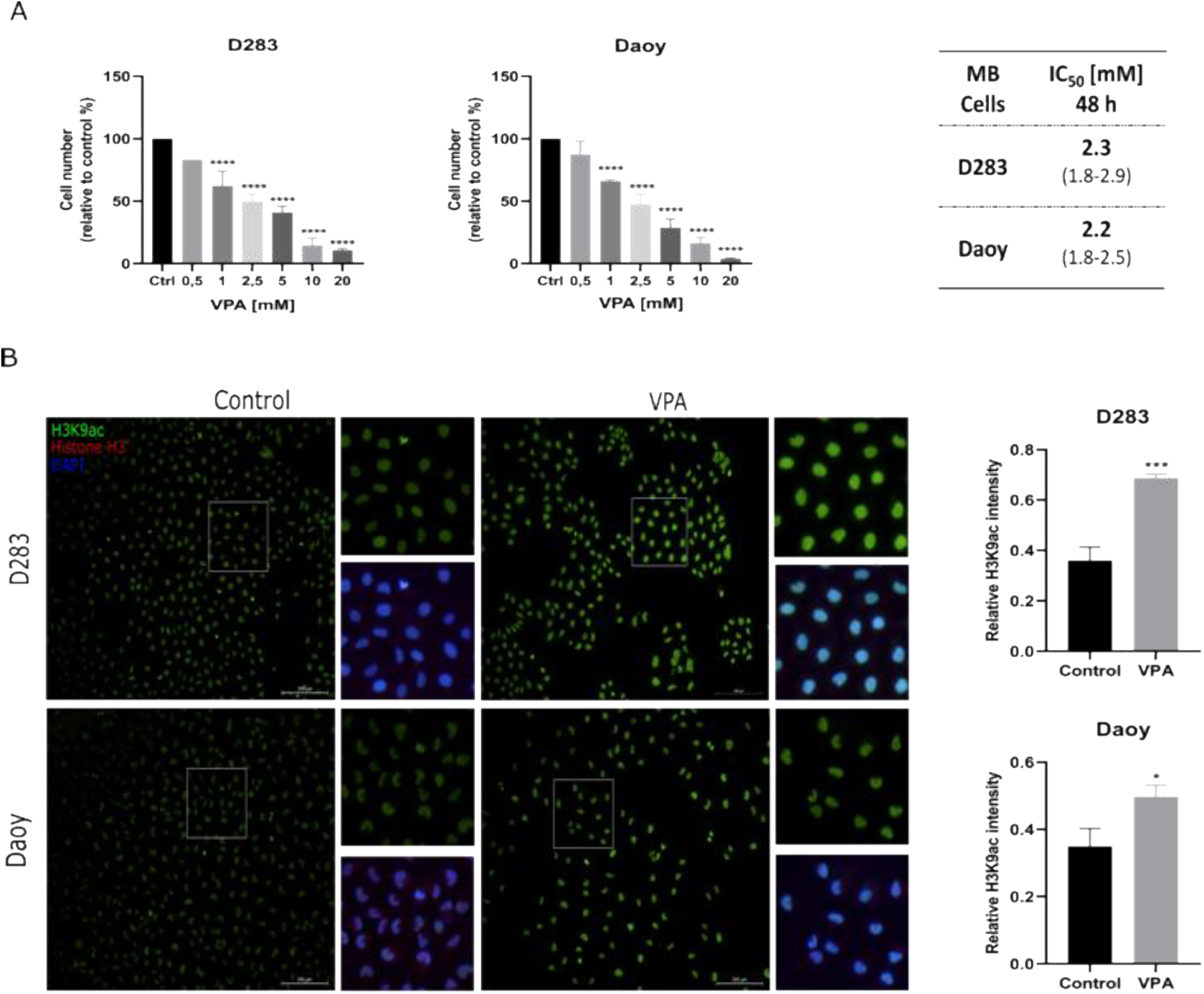
VPA reduces MB cell viability. (**A**) D283 and Daoy human MB cells were treated with a range of VPA concentrations (0.5, 1.0, 2.5, 5.0, 10.0, and 20.0 mM) for 48 h, and cell viability was measured by a trypan exclusion assay. IC_50_ concentrations of VPA for MB cells with a 95% confidence interval (CI) were 2.3 mM for D283 and 2.2 mM for Daoy cells. (**B**) Immuno- fluorescence assay using antibodies against H3K9ac, histone H3, and nuclei marker (DAPI) in the MB control and VPA-treated cells. The IC_50_ concentration of VPA (2.3 mM for D283 and 2.2 mM for Daoy) was used. Semi-quantification of the H3K9ac signal intensity relative to DAPI is shown. Results are the mean ± SD of three independent experiments; * *p* < 0.05, *** p < 0.001 and **** *p* < 0.0001 compared to the controls.

### 3.2. VPA Induces Cell Cycle Arrest Accompanied by Decreased MYC Expression in MB Cells

We measured the cell cycle distribution in MB cells after VPA exposure. Again, consistently with the previous study by Li et al. [29], VPA induced G1 arrest in the Daoy cells (*p* < 0.05) (Figure 2A). To better understand how VPA influences cell cycle progression, we examined the expression of *CDKN1A*, the gene that encodes cell cycle regulator protein p21, and *MYC*, which encodes the MYC transcription factor. *CDKN1A* and *MYC* levels were not significantly altered in the D283 cells. In the Daoy cells, VPA was able to increase the transcriptional levels of *CDKN1A* (4.2-fold, *p* < 0.05) while reducing *MYC* (0.7-fold, *p* < 0.0001) (Figure 2B). In addition, the p21 protein levels were slightly reduced in the VPA-treated D283 cells (0.78- fold, *p* < 0.05) and significantly increased in the Daoy cells (1.97-fold, *p* < 0.05) (Figure 2C).

**Figure 2.**
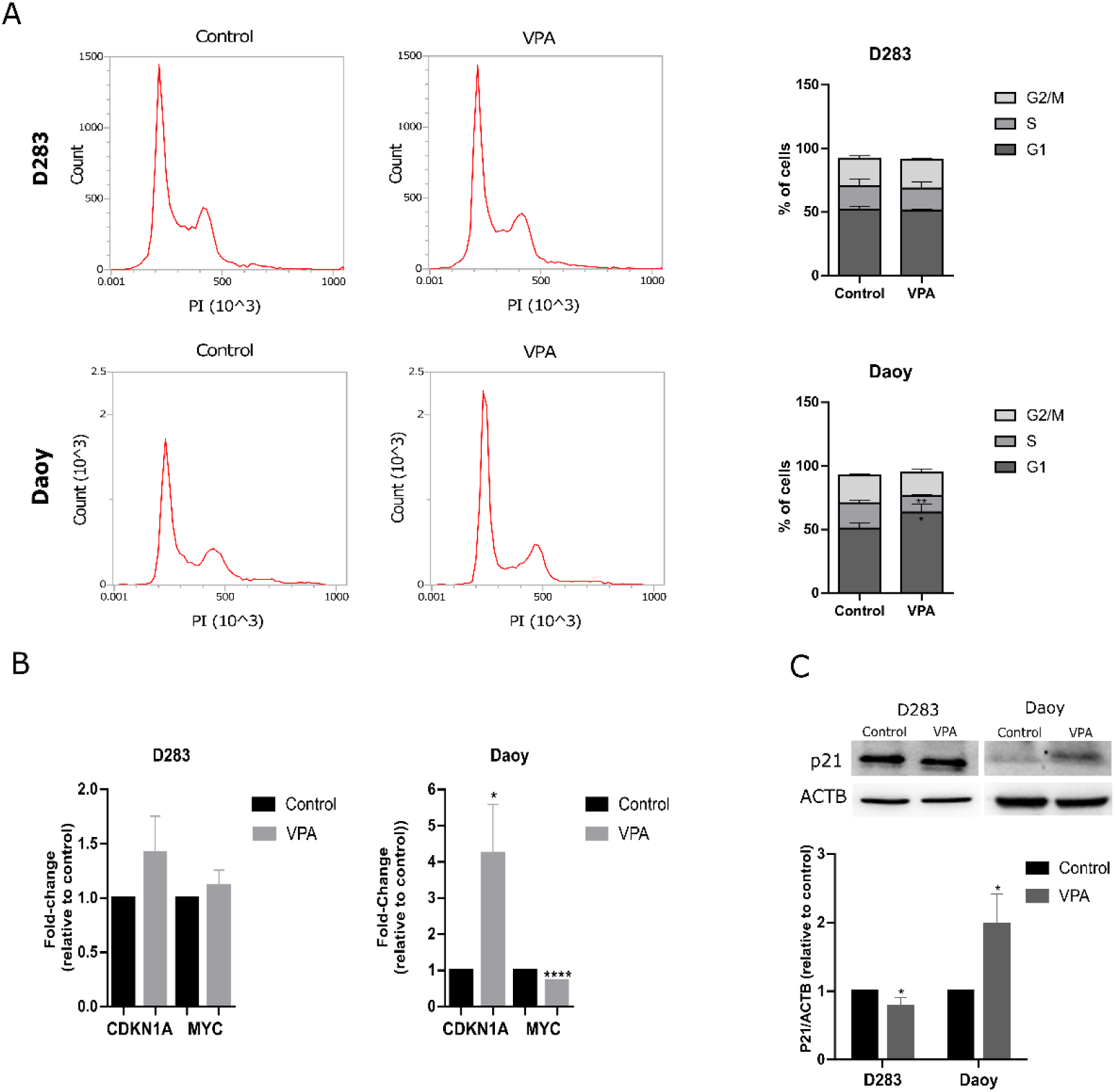
VPA leads to cell cycle arrest in MB cells. (**A**) Cell cycle analysis of the D283 and Daoy cells treated with VPA and controls. (**B**) Relative mRNA levels of *CDKN1A* and *MYC* in MB cells were assessed using RT-qPCR. (**C**) Western blot analysis of the p21 protein in MB cells after VPA exposure. The relative densitometric unit (RDU) analysis normalized by β-actin and corrected by the control is shown. All experiments were conducted using the IC_50_ concentration of VPA (2.3 mM for D283 and 2.2 mM for Daoy) for 48 h. Results are the mean ± SD of three independent experiments; * *p* < 0.05, ** *p* < 0.01 and **** *p* < 0.0001 compared to the controls.

### 3.3. VPA Reduces Formation of MB Neurospheres Enriched in Stemness Genes

Neurosphere formation assays are widely used to expand MB CSCs [14,22,33,35]. First, we verified the expression of stemness genes SOX2, *NES,* and *PRTG* in the D283 and Daoy neurospheres compared to the monolayer cells. After 7 days of culturing cells in an appropriate medium for the expansion of MB neurospheres, the D283 and Daoy-derived neurospheres showed an increase in the transcriptional levels of both stemness genes (1.9-fold, *p* < 0.01 for *SOX2* and 3.2-fold, *p* < 0.01 for *NES* in D283; 27-fold, *p* < 0.01 for *SOX2* and 64.4-fold, *p* < 0.0001 for *NES* in Daoy). *PRTG* transcriptional levels were upregulated in the D283 neurospheres and downregulated in the Daoy neurospheres (1.6-fold, *p* < 0.05 in D283 spheres and 0.5-fold, *p* < 0.01 for the Daoy neurospheres). Overall, these results suggest that the MB neurosphere assay enriched the stemness markers (Figure 3A,B).

**Figure 3.**
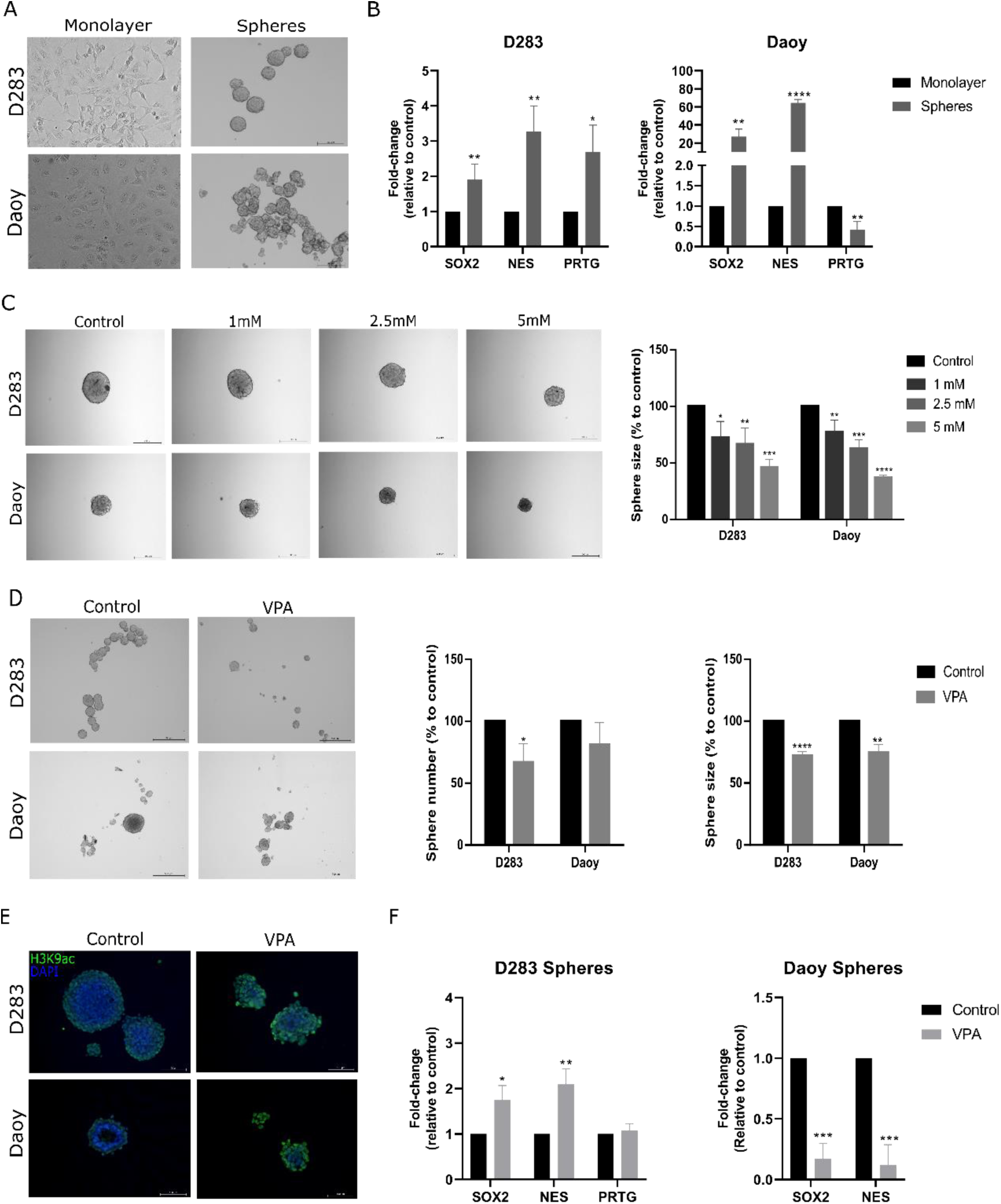
VPA impairs the expansion of MB neurospheres enriched with stemness marker genes. (**A**) Representative images of D283 and Daoy MB cells and derived neurospheres. (**B**) Relative mRNA levels of *SOX2, NES,* and *PRTG* in the monolayer cells and neurospheres were verified using RT-qPCR. (**C**) VPA effects on MB neurosphere formation after 5 days of drug exposure. Semi-quantitative analysis of neurosphere size in the VPA-treated and control neurospheres is shown. (**D**) After 5 days of MB neurosphere growth, VPA was added and the neurosphere size and number were evaluated after 48 h. Semi-quantitative analyses of the neurosphere size and number in the VPA-treated and control neurospheres are shown. Images were taken in an inverted microscope with 5 X magnification and the scale bar corresponded to 500 μm. (**E**) An immunofluorescence assay against H3K9ac, histone H3, and nuclei marker (DAPI) was performed in the VPA-treated and control neurospheres. Fluorescent images were taken with an inverted microscope with 20 X magnification and the scale bar corresponded to 100 μm. (**F**) Relative mRNA levels of *SOX2*, *NES,* and *PRTG* in the control and VPA-treated neurospheres were verified using RT-qPCR. Experiments were conducted using the IC_50_ concentration (2.3 mM for D283 and 2.2 mM for Daoy) of VPA. Results represent the mean ± SD of three independent experiments; * *p* < 0.05; ** *p* < 0.01; *** *p* < 0.001; and **** *p* < 0.0001 compared to the controls.

To elucidate whether VPA could influence MB neurosphere formation, we measured the neurosphere size after growth in the presence of VPA (1.0, 2.5, or 5.0 mM). All concentrations tested significantly reduced the neurosphere size after 5 days of VPA exposure compared to the controls in both cell lines (Figure 3C). We also examined whether VPA could reduce the size and number of neurospheres. After 5 days of neurosphere formation, neurospheres were treated with VPA (2.3 mM for D283 and 2.2 mM for Daoy) for 48 h. VPA reduced the D283 neurosphere size by 27% (*p* < 0.0001 compared to the controls) and number by 33% (*p* < 0.05 compared to the controls). In Daoy, VPA reduced the neurosphere size by 25.5% (*p* < 0.01 compared to the controls; Figure 3D). The immunofluorescence assay in MB neurospheres indicated that VPA at concentrations that impair MB neurosphere viability is also able to enhance H3K9ac (Figure 3E). After VPA treatment, the D283 neurospheres showed an increase in *SOX2* and *NES* (1.7-fold, *p* < 0.05 for *SOX2* and 2.0-fold, *p* < 0.01 for *NES*) whereas the *PRTG* levels were not significantly changed. In the Daoy spheres, the reduction in neurosphere viability was accompanied by a decrease in the *SOX2* and *NES* levels (0.2-fold, *p* < 0.001 for *SOX2* and 0.2-fold, *p* < 0.001 for *NES*) (Figure 3F).

### 3.4. VPA Reduces MYC and Increases TP53 Levels in MB Neurospheres

VPA increased the *CDKN1A* levels in the MB neurospheres (1.9-fold, *p* < 0.05 in D283 and 7-fold, *p* < 0.0001 in the Daoy-derived neurospheres) and decreased the *MYC* levels in both the D283 (0.7-fold, *p* < 0.05) and Daoy neurospheres (0.2-fold, *p* < 0.01) (Figure 4A). In the D283 neurospheres, VPA also increased the *TP53* levels (20-fold, *p* < 0.01) (Figure 4B) Moreover, the ChIP-qPCR results indicated that *TP53* may be epigenetically modulated by VPA in the D283 neurospheres (Figure 4C).

**Figure 4.**
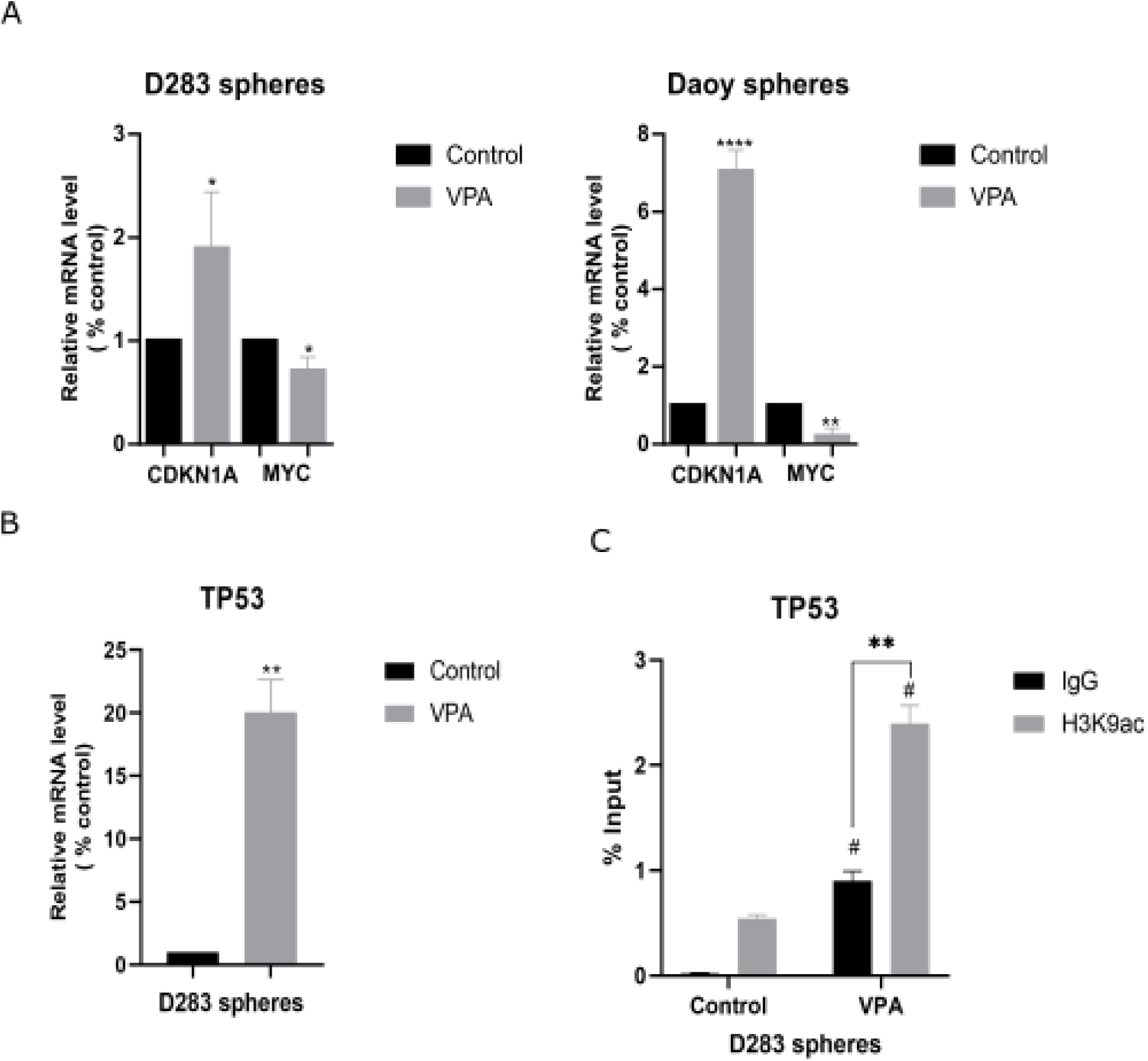
VPA increases *CDKN1A* and *TP53* while reducing *MYC* in MB-derived neurospheres. (**A**) Relative mRNA levels of *CDKN1A* and *MYC* in the D283- and Daoy-derived MB neurospheres were verified using RT-qPCR. (**B**) Relative mRNA levels of *TP53* in the D283- derived MB neurospheres were verified using RT-qPCR. (**C**) ChIP-qPCR for H3K9ac occupancy of the promoter region of *TP53* in the D283-derived neurospheres. Experiments were conducted using the IC_50_ concentration (2.3 mM for D283 and 2.2 mM for Daoy) of VPA for 48 h. Results are the mean ± SD of three independent experiments; * *p* < 0.05, ** *p* < 0.01, and **** *p* < 0.0001 compared to the controls. For the ChIP results * *p* < 0.05 and ** *p* < 0.01 comparing the VPA-treated cells versus the controls immunoprecipitated with anti-IgG or anti- H3K9ac; # *p* < 0.05 comparing the samples immunoprecipitated with the same antibody (IgG versus H3K9ac) between the VPA-treated and control groups.

### 3.5. Neuronal Differentiation and Changes in Stemness Markers Promoted by VPA in MB Cells

Exposure to VPA was accompanied by morphological changes consistent with differentiation in the MB cells (Figure 5A). We measured the transcriptional levels of neuronal differentiation markers *TUBB3* and *ENO2* and found that VPA increased the expression of both markers in the D283 (5.6-fold, *p* < 0.01 for *TUBB3* and 2.5-fold, *p* < 0.01 for *ENO2*) and Daoy cells (1.4-fold, *p* < 0.01 for *TUBB3* and 2-fold, *p* < 0.01 in *ENO2*; Figure 5B). In the MB-derived neurospheres, VPA also increased both markers (D283-derived neurospheres, 2.9-fold, *p* < 0.0001 for *TUBB3* and 2-fold, *p* < 0.001 for *ENO2*; Daoy-derived neurospheres, 7.1-fold, *p* < 0.01 for *TUBB3* and 13.1-fold, *p* < 0.01 for *ENO2*; Figure 5D). Immunofluorescence was used to detect the beta III tubulin protein levels after VPA exposure in the MB monolayer cells and neurospheres (Figure 5C,E).

**Figure 5.**
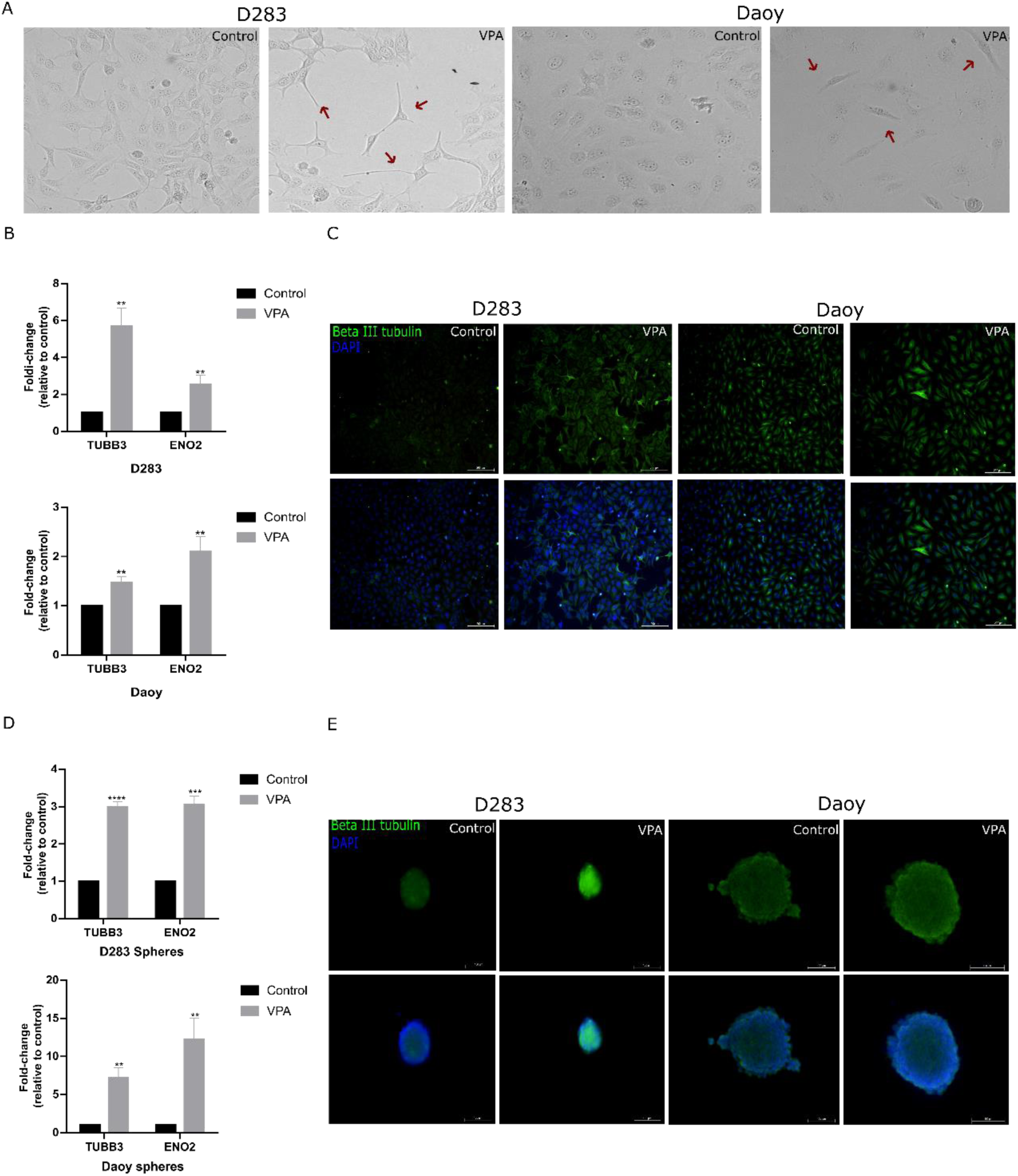
VPA effects on features of neuronal differentiation in MB cells. (**A**) Representative images of morphological changes in the VPA-treated and control D283 and Daoy MB cells. (**B**) Relative mRNA levels of *TUBB3* and *ENO2* in the MB cells were verified using RT-qPCR. (**C**) Immunofluorescence assay against neuronal differentiation marker protein beta III tubulin and nuclei marker (DAPI). (**D**) Relative mRNA levels of *TUBB3* and *ENO2* in the VPA-treated and control neurospheres derived from either D283 or Daoy cells were measured using RT-qPCR. (**E**) Immunofluorescence assay against neuronal differentiation marker beta III tubulin and nuclei marker (DAPI) in the VPA-treated and control neurospheres. The IC_50_ concentration of VPA (2.3 mM for D283 and 2.2 mM for Daoy) was used for treatment. Cells were exposed to VPA for 48 h. Fluorescent images were taken with an inverted microscope with a magnification of 10 X (monolayer) or 20 X (neurospheres). Results are the mean ± SD of three independent experiments; ** *p* < 0.01, *** *p* < 0.001, and **** *p* < 0.0001 compared to the controls.

Analysis of MB tumors from patients for the expression of VPA-regulated stemness-regulating genes showed that the expression of *MYC* in group 3 and WNT MB was significantly higher when compared to the group 4 and SHH molecular subgroups (Supplementary Figure S1A). High levels of *MYC* transcripts were associated with better prognosis as assessed by significantly longer OS in patients with SHH MB (Supplementary Figure S1B) but shorter OS in group 3 and group 4 MB (Supplementary Figure S1C,D). Thus, higher *MYC* expression may indicate poorer prognosis in more aggressive subgroups of MB. In addition, *NES* transcription was significantly higher in WNT MB tumors compared to all other subgroups, and was also higher in group 4 compared to group 3 and SHH MB (Supplementary Figure S2A). Higher *NES* expression was associated with worse prognosis, indicated by shorter OS only in patients with SHH MB (Supplementary Figure S2B).

## 4. Discussion

MB is a pediatric tumor type that presents high frequencies of mutations in genes encoding regulators of epigenetic processes, stemness, and differentiation [16]. The resulting cellular dysregulation is also associated with the proliferation and function of MB CSCs [36]. Modulation of the histone acetylation landscape is considered a therapeutic alternative to target stemness pathways responsible for CSC maintenance [37]. VPA is a well-known anti-convulsant drug that acts at least partially as an HDAC inhibitor [20]. Previous experimental studies on MB indicate that VPA induces antitumor effects in vitro and in vivo, alters pathways related to cell cycle progression, senescence, and apoptosis, and prolongs survival rates [29,38–43]. Here, we report that VPA displays inhibitory effects on the viability of MB cell lines representative of group 3 and SHH molecular subgroups as well as in neurospheres enriched with putative MB CSCs. The effects of VPA may be related to changes in cell cycle progression, stemness maintenance, and differentiation.

Our results indicate that VPA selectively induces cell cycle arrest in Daoy cells and affects molecular markers associated with cell cycle arrest in both D283- and Daoy-derived neurospheres, possibly by modulating p21 and *MYC*. These findings largely confirm those previously reported by Li et al. [29]. p21 is a negative regulator of the cell cycle, and *MYC* is a known modulator of p21 expression due to its ability to bind to the promoter region of *CDKN1A* and thus repress transcription [44,45]. VPA leads to *MYC* downregulation, which could explain the increase in p21 in Daoy cells. Data available in the Human Protein Atlas database (http://pro-teinatlas.org) confirmed different *MYC* expression in the D283 and Daoy cells [46]. Within a putative CSC context, we found the downregulation of *MYC* in neurospheres derived from both cell lines as well as an increase in p21 transcriptional levels after VPA treatment. These findings support the possibility that the modulation of *MYC* is important in mediating cell cycle arrest.

We showed that VPA increases the *TP53* transcription levels in D283 neurospheres, possibly through mechanisms related to epigenetic modulation. *TP53* is a tumor suppressor gene estimated to be mutated in about 50% of all human cancer types, with events leading to *TP53* repression being major drivers of malignant transformation. The *TP53* product, the p53 protein, is a sequence-specific DNA binding protein, with its most well-known functions being the promotion of cell cycle arrest and apoptosis in response to various cellular stress stimuli. The cell cycle arrest response mediated by p53 depends on its direct transcriptional activation of *CDKN1A*/p21, culminating in the transcription repression of cell cycle genes [47–50]. p53 can also repress *MYC* expression in a p21-independent manner, which is necessary for efficient p53-mediated cell cycle arrest and differentiation [51]. In addition, p53 has been implicated in developmental and cell differentiation processes [52], acting as a negative regulator of proliferation and survival in NSCs, thus potentially suppressing tissue and CSC self-renewal [53]. Together, this evidence suggests that in the *TP53* wild-type, *MYC* overexpressing D283-derived neurospheres, p53-mediated mechanisms may contribute to the modulation of stemness features observed in response to VPA.

We found that VPA is capable of increasing the expression of neuronal differentiation markers. Histone acetylation is a crucial process regulating neural differentiation [54]. During nervous system development, VPA reduces cell proliferation and initiates differentiation by increasing the expression of genes including *NEUROG1*, *TUBB3*, and *MAP2*. VPA but not valpromide (an amide of VPA that does not inhibit HDACs) increases neuronal differentiation markers. Moreover, in NSCs from the rat embryo hippocampus, VPA causes an increase in H3K9ac active marks at the *NEUROG1* promoter, suggesting that epigenome alterations caused by VPA allow for the establishment of a more differentiated phenotype [55–57]. Given that MB has granule precursor and NSCs as possible cells of origin, it can be expected that VPA is able to induce morphological changes consistent with differentiation and promote the expression of neural differentiation markers [1, 2]. Once MB cells differentiate, they lose proliferative capacity and tumorigenic potential through processes dependent on epigenetic modulation [58].

SOX2 is associated with stemness regulation, self-renewal, pluripotency, and neural differentiation in embryonic stem cells (ESCs) [59,60]. In MB, SOX2 protein expression is critically involved in tumor development, especially in the SHH molecular subgroup [61]. GLI1/2 are downstream factors of SHH and positively regulate SOX2 by binding to its promoter, stimulating self-renewal and tumorigenesis [62]. In addition, SOX2-positive cells show lower sensitivity to chemotherapy agents [63]. Our results indicate that VPA can reduce the *SOX2* and *NES* transcriptional levels in Daoy neurospheres (*TP53*-mutated, representative of SHH) while increasing these markers in D283 neurospheres (*TP53* wild-type, group 3). *NES* is a progenitor marker regulated by SOX2 [64], suggesting that VPA also affects signaling components downstream of SOX2 [65].

Expression of the protogenin protein, encoded by the *PRTG* gene, is strong during a specific developmental interval in the embryonic neural tube, defining a transition stage of neurogenesis and blocked differentiation [66]. A recent study identified *PRTG*-expressing stem cells in a perivascular niche of the embryonic hindbrain that undergo oncogenic transformation to initiate group 3 MB [67]. Our finding that *PRTG* gene transcription was enhanced after neurosphere induction in D283 but not DAOY cells supports the view that protogenin acts as a specific driver of group 3 MB.

At least two genes influenced by VPA treatment in MB cells, *MYC* and *NES*, showed distinct patterns of expression among MB tumors belonging to different molecular subgroups. We found that higher *MYC* expression is associated with worse prognosis as indicated by shorter survival, which is consistent with previous evidence showing that group 3 MB tumors harbor *MYC* amplification [4,7,9,67], and patients with *MYC*-amplified MB have a poorer prognosis despite intensive treatment regimens [34,68]. Importantly, class I HDAC inhibitors are effective against *MYC*-amplified MB cells [69]. We also found that higher *NES* expression was associated with shorter OS in patients with SHH MB. A high content of Nestin, the protein encoded by the *NES* gene, is a feature of MB CSCs [70] and SHH-driven cancers [71]. Other factors related to stemness could be investigated in relation to modulation by HDACs in MB. For example, Krüppel-like factor 4 (KLF4) is one of factors used for the induction of pluripotent stem cells (iPS cells) from mouse embryonic or adult fibroblasts [72]. The expression of KLF4 in tumors is regulated by a range of epigenetic mechanisms including histone acetylation [73]. Loss of KLF4 expression both at the RNA and protein levels is found in over 40% of primary MB tumors, and the re-expression of KLF4 in D283 MB cells suppresses experimental MB growth [74].

Together with previous studies, our results suggest that the potential of VPA in MB therapy warrants further exploration. A case of the successful use of dose-dense neoadjuvant chemotherapy combined with VPA in a child with MB has been reported [75]. Our study, however, has several limitations. Cell line cultures are limited as disease models, and in vivo experiments using mice will be required to confirm and extend the present results and explore the dose–effects relationships. Additional MB cell lines could have been studied such as the *TP53* wild-type ONS76 line. The IC_50_ we found for the inhibition of MB viability was over three times higher than the plasma therapeutic concentration for VPA used in the clinical setting, and only 10 to 20% of the plasma concentration reached the cerebrospinal fluid (CSF). A concentration of only 0.4 mM of VPA is necessary for HDAC1 inhibition, which falls within the therapeutic range [76,77]. A requirement for very high (and potentially toxic) doses of VPA may thus be a challenge for its repurposing as an anticancer drug to treat MB.

## 5. Conclusions

In conclusion, our results indicate that VPA can influence MB viability and features related to cell cycle regulation, stemness, and differentiation, possibly by increasing histone acetylation, reducing *MYC*, and increasing *TP53*. A schematic summary of the key findings is shown in Figure 6. It is possible that VPA-induced inhibition of HDAC activity and the MYC–p21– SOX2 axis influence the development and aggressiveness of MB tumors.

**Figure 6.**
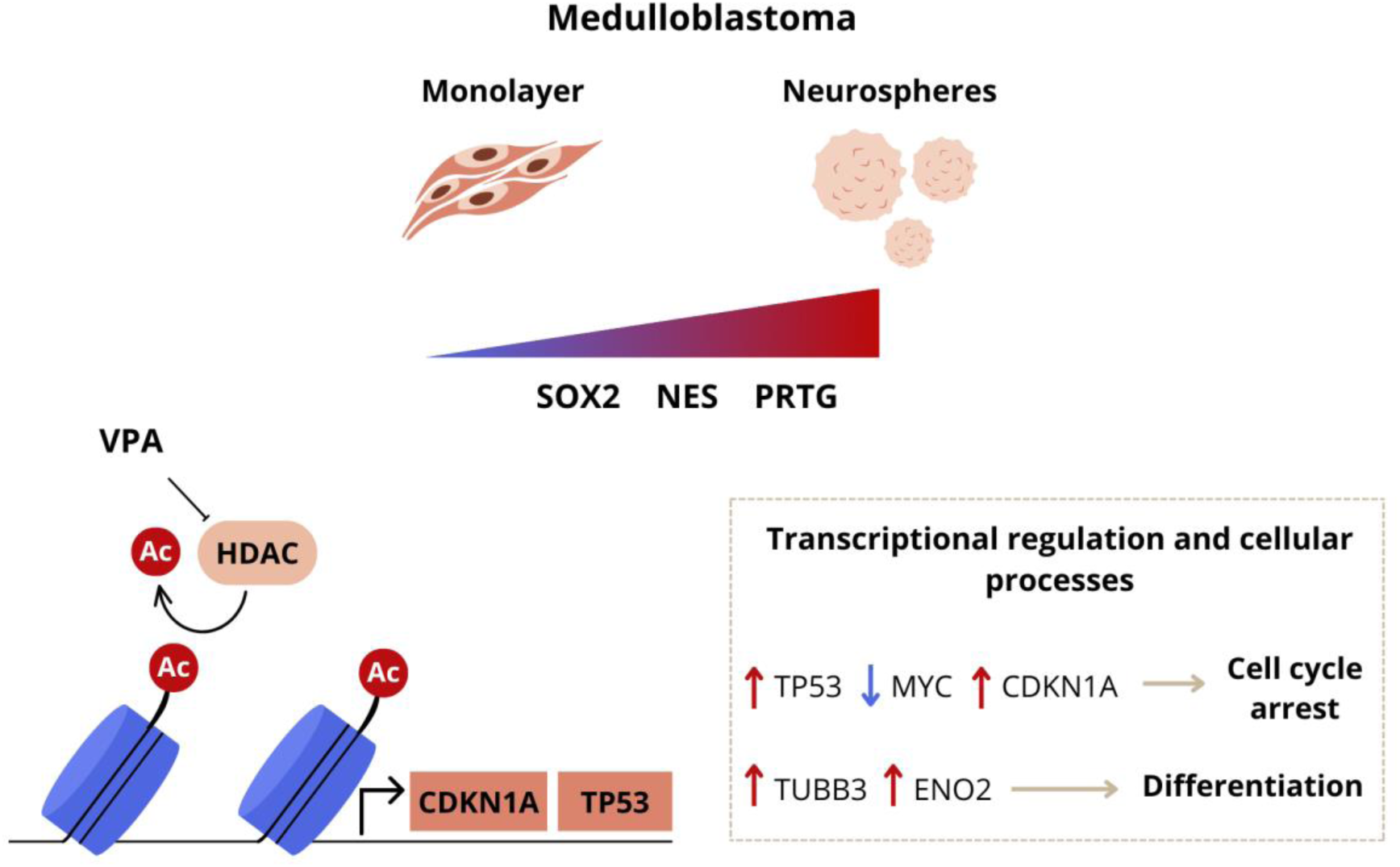
Schematic model highlighting selected aspects of VPA-induced molecular and functional effects in MB. Exposure to VPA leads to a reduction in MB cell cycle arrest and differentiation in the MB monolayer and neurospheres. VPA can increase *TP53* while reducing *MYC* expression, potentially inhibiting stemness and promoting features of differentiation.

## Supplementary Materials

The following supporting information can be downloaded: Supplementary Tables 1 and 2, Supplementary figures S1, S2, and S3.

## Author Contributions

Conceptualization, N.H.F., C.N., M.D.T., and R.R.; Methodology, N.H.F., A.L.H., J.V., M.D., M.A.C.F., C.N., V.R., C.B.d.F., A.T.B., M.d.C.J., M.D.T., and R.R.; Investigation, N.H.F., A.L.H., J.V., M.D., M.A.C.F., and M.d.C.J.; Resources, N.H.F., M.D., M.A.C.F., C.B.d.F., A.T.B., A.L.B., M.d.C.J., M.D.T., and R.R.; Data curation, N.H.F., A.L.H., J.V., M.D., M.A.C.F., and M.d.C.J.; Writing—original draft preparation, N.H.F. and R.R.; Writing—review and editing, N.H.F., A.L.H., J.V., M.D., M.A.C.F., C.N., V.R., C.B.d.F., A.T.B., A.L.B., L.J.G., M.d.C.J., M.D.T., and R.R.; Supervision, M.A.C.F., M.d.C.J., M.D.T., and R.R.; Project administration, N.H.F., C.B.d.F., A.T.B., A.L.B., M.d.C.J., and R.R.; Funding acquisition, M.A.C.F., C.N., V.R., C.B.d.F., A.T.B., A.L.B., L.J.G., M.d.C.J., M.D.T., and R.R. All authors have read and agreed to the published version of the manuscript.

## Funding

This work was supported by the National Council for Scientific and Technological Development (CNPq, MCTI, Brazil), grant numbers 305647/2019-9, 405608/2021-7, and 406484/2022-8 (INCT BioOncoPed) to R.R., 101566/2022-0 to A.L.H, and a scholarship to N.H.F., Ministry of Health/CNPq/FAPERGS PPSUS grant number 21/2551-0000114-3 to R.R. and L.G., the Children’s Cancer Institute (ICI), and Clinical Hospital (HCPA) institutional research fund grant numbers 2020-0069 and 2023-0021 to R.R. C.N. was supported by a William Donald Nash Brain Tumor Research Fellowship, Brain Tumor Foundation of Canada, and the Swifty Foundation. V.R. was supported by operating funds from the Canadian Institutes for Health Research, the Garron Family Cancer Center, the C.R. Younger Foundation, the Canadian Cancer Society Research Institute, Brain Canada, the Rally Foundation for Childhood Cancer, and a Canada Research Chair in Pediatric Neuro-Oncology. M.D.T. is a Cancer Prevention and Research Institute of Texas (CPRIT) Scholar in Cancer Research (CPRIT— RR220051) and the Cyvia and Melvyn Wolff Chair of Pediatric Neuro-Oncology, Texas Children’s Cancer and Hematology Center. M.D.T. is also supported by the National Institutes of Health (NIH) (R01NS106155, R01CA159859, and R01CA255369), the Pediatric Brain Tumor Foundation, the Terry Fox Research Institute, the Canadian Institutes of Health Research, the Cure Search Foundation, the Matthew Larson Foundation (IronMatt), b.r.a.i.n.child, Meagan’s Walk, the Swifty Foundation, the Brain Tumor Charity, Genome Canada, Genome BC, Genome Quebec, the Ontario Research Fund, Worldwide Cancer Research, the V Foundation for Cancer Research, the Ontario Institute for Cancer Research through funding provided by the Government of Ontario, a Canadian Cancer Society Research Institute Impact grant, a Cancer Research UK Brain Tumor Award, a Stand Up To Cancer (SU2C) St. Baldrick’s Pediatric Dream Team Translational Research grant (SU2C-AACR-DT1113), and SU2C Canada Cancer Stem Cell Dream Team Research Funding (SU2C-AACR-DT-19-15) provided by the Government of Canada through Genome Canada and the Canadian Institutes of Health Research, with supplementary support from the Ontario Institute for Cancer Research through funding provided by the Government of Ontario. M.D.T. was also supported by the Garron Family Chair in Childhood Cancer Research at the Hospital for Sick Children and the University of Toronto.

## Institutional Review Board Statement

This study was conducted following legal, regulatory, and institutional requirements and was approved by the appropriate institutional committees.

No animal experiments, biological samples obtained from patients, or clinical data were used in this study.

## Informed Consent Statement

Not applicable.

## Data Availability Statement

The dataset analyzed in this study is available in the Gene Expression Omnibus repository, https://www.ncbi.nlm.nih.gov/geo/query/acc.cgi?acc=GSE85217. Other data can be provided upon request.

## Conflicts of Interest

The authors declare no conflicts of interest related to the contents of this manuscript.

## Notes

https://www.ncbi.nlm.nih.gov/geo/query/acc.cgi?acc=GSE85217

### Competing Interest Statement

The authors have declared no competing interest.

### Summary of Updates

This version of the manuscript has been revised to correct minor errors and improve overall manuscript quality.

