## Supplementary information for "Modulation of Stemness and Differentiation Regulators by Valproic Acid in Medulloblastoma Neurospheres"

**Supplementary Table 1.** Forward and reverse primers used for RT-qPCR amplification.

| Gene | Primer Forward (5'-3') | Primer Reverse (5'-3') |
|---|---|---|
| *ACTB* | AAACTGGAACGGTGAAGGTG | AGAGAAGTGGGGTGGCTTTT |
| *CDKN1A* | ACTCTCAGGGTCGAAAACGG | CTTCCTGTGGGCGGATTAGG |
| *ENO2* | AGCCTCTACGGGCATCTATGA | TTCTCAGTCCCATCCAACTCC |
| *NES* | GATCGCTCAGGTCCTGGAAG | GGGGTCCTAGGGAATTGCAG |
| *MYC* | TACAACACCCGAGCAAGGAC | AGCTAACGTTGAGGGGCATC |
| *PTRG* | TGCATGCAAGATTCATCCCACCC | TGCAATACTCCTGTTGGTAGGGCA |
| *SOX2* | CAGCTCGCAGACCTACATGA | GGGAGGAAGAGGTAACCACAG |
| *TUBB3* | CTCAGGGGCCTTTGGACATC | CAGGCAGTCGCAGTTTTCAC |
| *TP53* | ACCTATGGAAACTACTTCCTGAAA | CTGGGAGCTTCATCTGGACC |

**Supplementary Table 2.** Primers targeting the promoter region of *TP53* in ChIP analysis.

| Gene | Primer Forward (5'-3') | Primer Reverse (5'-3') |
|---|---|---|
| *TP53* | TTTAGCGCCAGTCTT GAGCA | GTATCTACGGCACCA GGTCG |

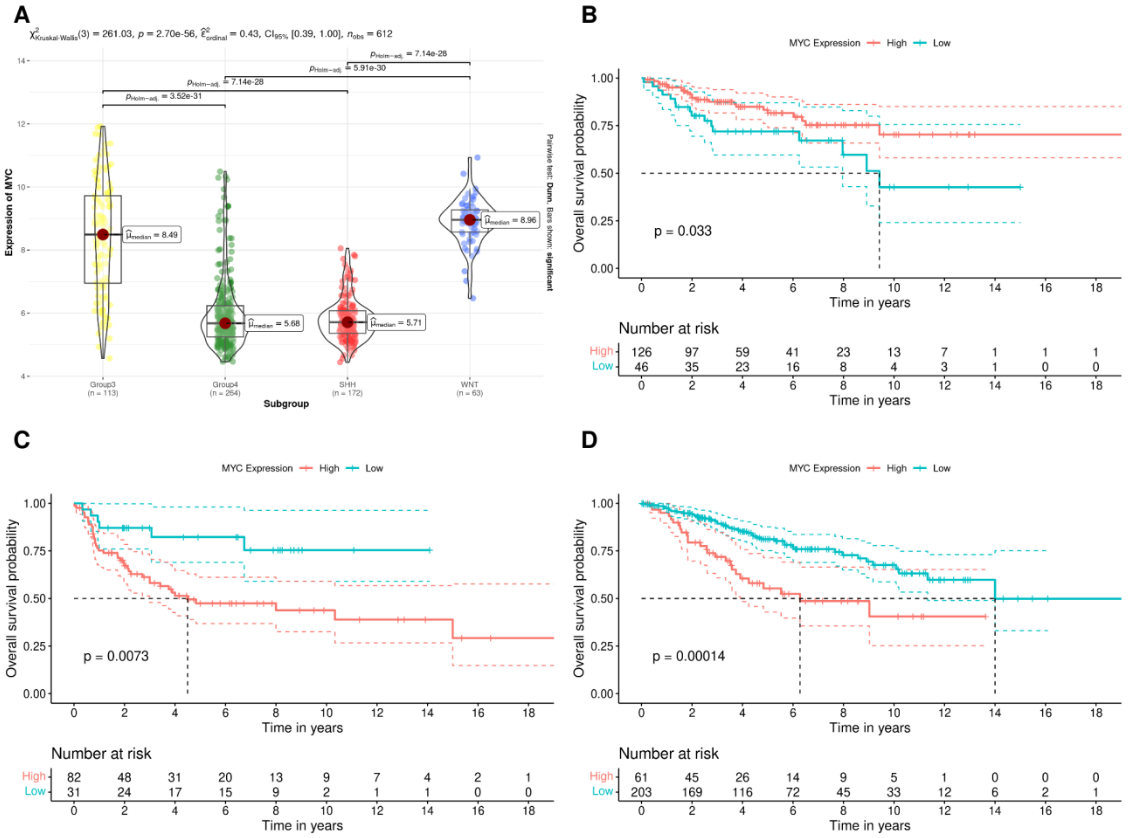


**Supplementary Figure S1.** *MYC* gene expression and association with patient prognosis across different molecular subgroups of human MB. **(A)** Transcript levels in group 3 ($n$ = 113), group 4 ($n$ = 264), SHH ($n$ = 172), and WNT ($n$ = 64). **(B-D)** OS of patients bearing MB tumors with high or low levels of *MYC* expression classified into different molecular subgroups. Analyses were performed as described in Materials and Methods; *P* values are indicated in the figure.

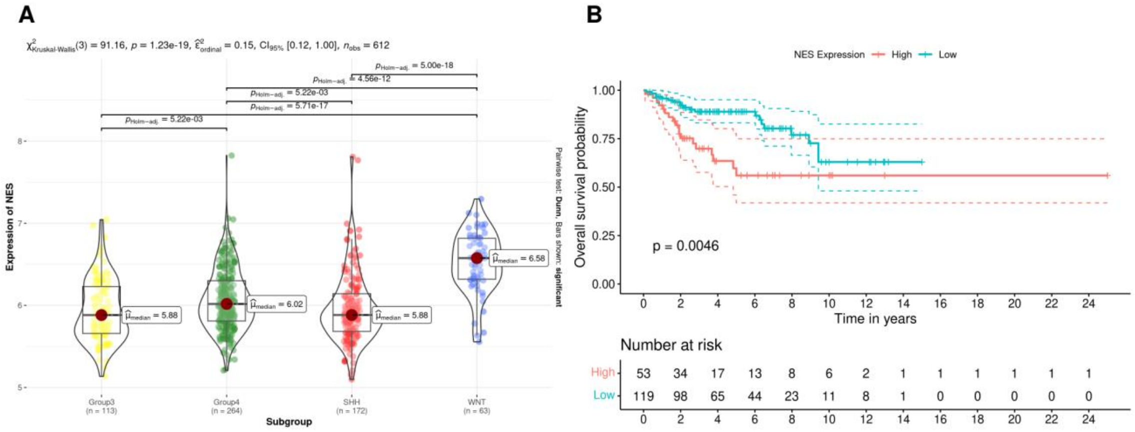


**Supplementary Figure S2.** *NES* gene expression across different molecular subgroups of human MB and association with prognosis in patients with SHH MB. (**A)** Transcript levels in group 3 ($n$ = 113), group 4 ($n$ = 264), SHH ($n$ = 172), and WNT ($n$ = 64). (**B)** OS of patients bearing SHH MB tumors with high or low levels of *NES* expression. Analyses were performed as described in Materials and Methods; *P* values are indicated in the figure.

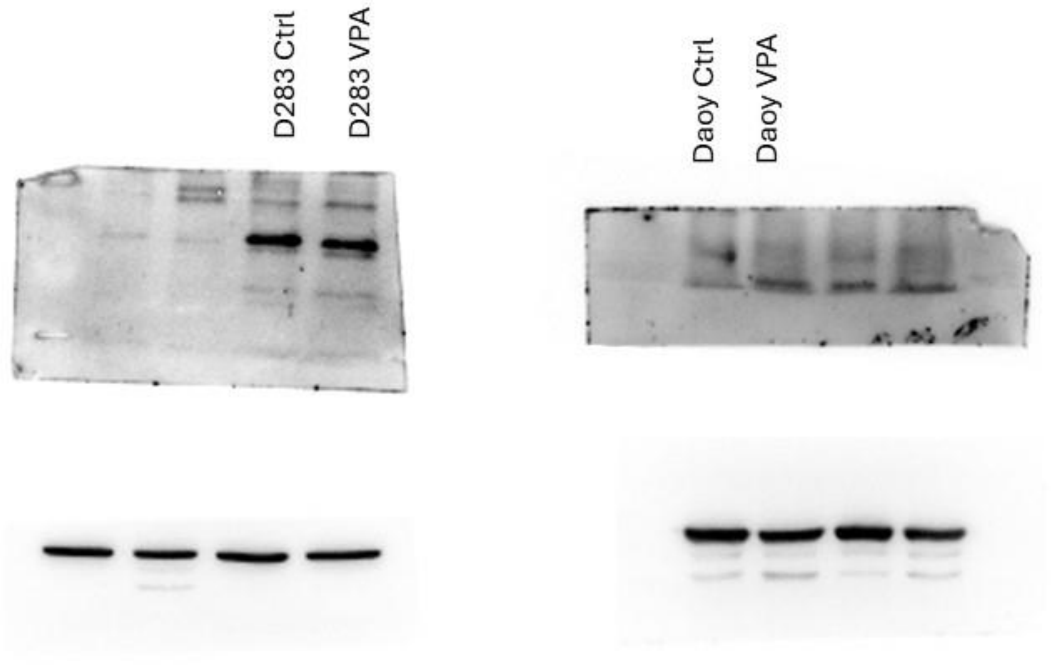


**Supplementary Figure S3.** Uncropped blots for the experiment shown in Figure 2C describing the Western blot analysis of p21 protein in D283 and Daoy MB cells.
